# Olive fruit fly rearing procedures affect the vertical transmission of the bacterial symbiont *Candidatus* Erwinia dacicola

**DOI:** 10.1101/367417

**Authors:** Patrizia Sacchetti, Roberta Pastorelli, Gaia Bigiotti, Roberto Guidi, Sara Ruschioni, Carlo Viti, Antonio Belcari

## Abstract

Background: The symbiosis between the olive fruit fly, *Bactrocera oleae*, and *Candidatus* Erwinia dacicola has been demonstrated as essential for the fly’s larval development and adult physiology. The mass rearing of the olive fruit fly has been hindered by several issues, including problems which could be related to the lack of the symbiont, presumably due to preservatives and antibiotics currently used in the laboratory. To better understand the mechanisms underlying symbiont removal or loss during the rearing of lab colonies of the olive fruit fly, we performed experiments that focused on bacterial transfer from wild female flies to their eggs. In this research, eggs laid by wild females were treated with propionic acid solution, which is often used as an antifungal agent, a mixture of sodium hypochlorite and Triton X, or water (as a control). The presence of the bacterial symbiont on eggs was evaluated by real-time PCR and scanning electron microscopy.

Results: DGGE analysis showed a clear band with the same migration behavior present in all DGGE profiles but with a decreasing intensity. Molecular analyses performed by real-time PCR showed a significant reduction in *Ca*. E. dacicola abundance in eggs treated with propionic acid solution or a mixture of sodium hypochlorite and Triton X compared to those treated with water. In addition, the removal of bacteria from the surfaces of treated eggs was highlighted by scanning electron microscopy.

Conclusions: The results clearly indicate how the first phases of the colony-establishment process are important in maintaining the symbiont load in laboratory populations and suggest that the use of products with antimicrobial activity should be avoided. The results also suggest that alternative rearing procedures for the olive fruit fly should be investigated.

## Background

Insects display a great variety of symbiotic relationships with microorganisms that allow them to exploit almost every substrate as food source and to colonize any habitat on earth. Such microorganisms comprise viruses as well as bacteria, fungi, protozoa and multicellular symbionts [1]. The relationships between insects and microorganisms range from clear mutualism to relationships involving unbalanced benefits or costs to one member up to pathogenesis [2, 3]. Moreover, insect symbioses can vary from temporary associations to long-life obligate partnerships and from external, loose coalitions to very close alliances [2]. The microorganisms involved can be found in the environment, growing outside the insect’s body, or they can be harbored within the body cavity in specialized cells or organs (extracellular or intracellular endosymbionts) and transmitted through successive generations, typically via vertical transmission from mother to progeny (maternal inheritance) [4]. The manifold and intricate functions played by the wide assortment of microorganisms have not been fully studied in detail, and only some metabolic interactions are fully understood [5]. Regardless, the crucial roles played by symbionts in the survival and evolution of their insect partners have been repeatedly demonstrated, and different mechanisms of transmission through host populations and generations have evolved [2].

The nonpathogenic bacterial symbionts of insects have been classified as ranging from primary, ancient obligate symbionts that are restricted to specialized cells (bacteriomes) and are necessary for the host to secondary, recent facultative symbionts that are located in insect organs and are non-essential for insect survival [1, 6]. The transmission of primary symbionts (P-symbiont) in plant-feeding insects has been investigated in detail in aphids [7, 8], various sucking insects [9, 10, 11] and beetles [12, 13]. Bacterial P-symbionts are transferred vertically to offspring through contamination of the egg surface, deposition of bacterial capsules on eggs, or consumption of the mother’s excrement or through transovarial transmission; that is, direct penetration of the female germ cells [11]. Maternal inheritance is also the typical transmission route for secondary symbionts, although there is substantial evidence of horizontal transmission as well as rare paternal transmission [14, 15].

Similarly to sucking insects, Tephritid fruit flies display many types of symbiotic associations involving both intracellular (e.g. Wolbachia), and extracellular symbionts. Lauzon [16] critically reviewed this topic, commenting on known features and highlighting important issues with possible practical consequences for insect pest control. Many tephritids are insect pests of economic importance, causing damage to agricultural crops in tropical, subtropical and temperate areas [17]. By studying the relationships of fruit fly species with symbiotic bacteria, new control strategies might be developed and established [18]. During the last decade, research on the symbiotic relationships of fruit flies has often focused on potential pest control applications. Moreover, following Lauzon’s review [16], research on this topic was greatly increased by the advent of molecular techniques, which enabled the investigation of uncultivable bacteria and thus the identification of previously unknown or misidentified microorganisms.

An example of a symbiotic relationship that was clarified via molecular techniques is that between the olive fruit fly, *Bactrocera oleae* (Rossi), which is the major insect pest of olive crops in countries where it occurs, and the bacterium *Candidatus* Erwinia dacicola, which was named in 2005 [19]. This symbiosis was the first one involving tephritids to be described, discovered at the beginning of the twentieth century, although the bacterium was erroneously identified as *Pseudomonas savastanoi*, the agent of olive knot disease. Relying only on microscopic observations, Petri [20, 21] carefully described a specialized foregut organ that harbored the symbiont (a cephalic evagination later named “oesophageal bulb”) as well as female hindgut pockets from which bacteria were released to be deposited on the egg surfaces and transmitted to the next generation. Since Petri’s investigations, several authors have increased knowledge on the olive fruit fly and bacterium symbiosis, providing indirect evidence of the essential role of the symbiont for the insect’s survival (see the reviews by Drew and Lloyd, [22], and Lauzon, [16]). However, there were no major findings until the discovery of PCR amplification and 16S rRNA gene sequencing techniques which have significant improved our knowledge on olive fruit fly symbiotic associations.

To summarize recent findings, we know that *Ca*. E. dacicola is an unculturable bacterium that belongs to the Enterobacteriaceae family of gammaproteobacteria [19]. This bacterium is considered an obligate symbiont (P-symbiont) that coevolved with its host *B. oleae* wherein it dwells extracellularly inside the adult gut (in the oesophageal bulb, crop, midgut and female rectal pockets) and the larval midgut (gastric caeca) [19, 23]; it also lives intracellularly inside epithelial cells of the larval midgut [23]. *Ca*. E. dacicola forms bacteriomes in the larval gut, whereas in adults, it typically develops biofilms that line the inner surfaces of organs or fills the lumen of different organs with abundant free bacterial masses [23, 24]. The species occurs as two different haplotypes in Italian populations of *B. oleae* [25, 26]. Regarding its roles in host physiology, the symbiont is essential for larvae, allowing them to feed on olives, mainly when they are unripe, and neutralizing the negative effects of the phenolic compound oleuropein [27]. Moreover, *Ca*. E. dacicola is necessary for adults of the olive fruit fly as it metabolizes complex nitrogen compounds and supplies growth factors that can promote fly survival and reproduction in food-inadequate habitats such as olive orchards [28, 29].

According to the observations by Petri [21], the symbiont is vertically transmitted to the progeny: When eggs exit the oviduct, they pass through the terminal rectal tract, where the rectal sacs open and bacterial masses are deposited onto the eggs’ surfaces. Then, at eclosion, larvae ingest bacteria breaking through the micropylar pole. This hypothesized mechanism of transmission is supported by ultrastructural investigations using SEM and TEM [23, 30], that show the presence of abundant bacteria stored in rectal evaginations in association with the genital and anal openings.

Having established the importance of *Ca*. E. dacicola for the regular development and adult fitness of the olive fruit fly, we can understand how the symbiotic relationship might be manipulated to improve the control of this pest. A few years ago, Estes and colleagues [31] reviewed knowledge on the possible application of the Sterile Insect Technique (SIT) for the olive fruit fly, highlighting critical issues, possible improvements and future directions. They emphasized the necessity of understanding the interactions between the insect pest and its symbiont in wild populations as well as the insect’s interactions with different bacteria in laboratory colonies. In nature, *B. oleae* larvae develop only in olives; a group of Greek scientists devoted more than 20 years to developing an artificial substrate suitable for its mass rearing [31, 32]. The symbiont *Ca*. E. dacicola has never been retrieved from lab-reared olive flies, which appear to be associated with a variety of bacteria, typically species that are common in colonies of lab-reared insects [23, 33, 34]. It is likely that the presence of the symbiont is prevented by the usage of preservatives and antibiotics that are typically added to larval and/or adult diets [32]. Moreover, the yield and quality of mass-reared olive fruit flies, in term of fitness and behavior, have yet to reach satisfactory levels [35, 36]. Therefore, only a few pilot trials of SIT application have been attempted, with unsatisfactory results [37, 38, 39]. The first step toward developing feasible SIT programs is to reevaluate the mass rearing of the olive fruit fly, taking into consideration what we know about its symbiont. We believe that two approaches should be pursued: a) supply lab flies with diet-enriched transient bacteria to potentially replace the role played by the natural symbiont *Ca*. E. dacicola and b) begin the colonization process anew from wild symbiotic olive fruit flies while avoiding symbiont-removing or symbiont-suppressing procedures in the rearing protocol.

The first approach was recently initiated with promising results [40], while the second approach has to be initiated, although the rearing of wild olive fruit flies on an antibiotic-free diet for eight generations has been attempted [41].

The present study is part of a long-term research program addressing the multiple relationships between *B. oleae* and bacteria and aimed at identifying target points that might be used to develop new control strategies. To evaluate the effects of commonly used procedures to rear olive fruit flies in the laboratory on *Ca*. E. dacicola, we assessed the effects of disinfectants that are used for handling eggs, which is the first step in both small-scale and large-scale rearing efforts, through PCR amplification-denaturing gradient gel electrophoresis (PCR-DGGE), quantitative real-time PCR and Scanning Electron Microscopy (SEM). In addition, by evaluating the impacts of germicides, we ascertained the transmission mechanism of *Ca*. E. dacicola from wild olive fruit fly females to their progeny reared in laboratory.

## Methods

### Insects

The adults of wild olive flies used in this study developed from pupae that had been collected from infested drupes in several olive orchards in Vaccarizzo Albanese (Cosenza; Italy). Flies had been housed in plastic cages (BugDorm-1, MegaView Science, Taiwan), with approximately 800 flies per cage, supplied with sugar and water, and maintained at room temperature (18–20 °C). To enhance egg production, flies were transferred into a conditioned rearing room with conditions of 25±2 °C, 60±10% RH and a 16:8 (L:D) photoperiod and supplied a diet of sugar, hydrolyzed enzymatic yeast (ICN Biomedicals) and egg yolk (40:10:3).

### Egg collection

The eggs of wild flies were collected using wax domes that had been washed previously with 2% hypochlorite solution and then rinsed twice with deionized water. The domes were inserted into the bottom of tissue culture dishes (35/10 mm) containing approximately 3 mL of deionized water. These measures were taken to minimize the occurrence of contaminating bacteria and prevent egg dehydration and subsequent shrinkage. The domes were placed inside the adults’ cage and left there for 24 hours. Eggs were then collected by washing the internal surface of the domes with sterile deionized water under a laminar flow hood and sieving with a sterile cloth; the eggs were then placed in a sterile beaker. Finally, the eggs were collected with a sterile micropipette and transferred to three different sterile containers.

The three containers contained the following treatments, respectively: a 0.3% propionic acid solution (PA) (pH=2.82±0.03) commonly used as disinfectant in rearing procedures of the olive fruit fly [32]; a mixture (1:1) of 1% sodium hypochlorite + 0.1% Triton X (SHTX) previously used to externally sterilize all of the developmental stages of the olive fruit fly by Estes *et al*. [42]; and sterile water as a control (C). All the eggs were vortexed for 30 s, and then the eggs of the treatments PA and SHTX were rinsed twice in deionized sterile water (in order to remove treatment residues which would have hampered DNA extraction). Eggs of each group (PAE, SHTXE, CE, respectively) were designated for microbiological analyses as well as for morphological observations or larval development. Egg collection was performed four times during the experiment, each time from a different cage.

In addition, and in order to evaluate the bacterial titer of the water or rinse water where eggs were taken from, liquid samples were also collected for further molecular analysis: egg collection water of the control treatment (CW), the second rinse water after 0.3% propionic acid treatment (PAW) and the second rinse water after SHTX treatment (SHTXW).

An explanatory list of the samples analyzed in the experiment is summarized in Table 1.

**Table 1.** Explanatory legend of samples analyzed in the egg treatment experiment

| Sample description | Sample name |
| --- | --- |
| Eggs washed with water (control) | CE |
| Eggs treated with 0.3% propionic acid | PAE |
| Eggs treated with a mixture (1:1) of 1% sodium hypochlorite + 0.1% TritonX | SHTXE |
| Water from control eggs | CW |
| Second rinse water after treatment with PA | PAW |
| Second rinse water after treatment with SHTX | SHTXW |

### Progeny development

This experiment was carried out in the same conditioned rearing room described above. Eggs intended for larval development were spread over a black fabric disk soaked in water and positioned in a Petri dish. After 48 hours, the hatched and unhatched eggs were counted. Each group of larvae from the different egg treatments (CE, PAE, SHTXE) was transferred to a cellulose-based artificial diet [32] until pupation. Then, the pupae were collected and placed in vials for adult emergence. Newly emerged adults were singly placed in small cages and fed with water and sugar until they were 15 days old, when they were dissected for bacterial DNA extraction.

### DNA extraction from eggs and DGGE analysis

Ten eggs per treatment were sampled under the stereomicroscope and transferred into a 1.5 mL tube containing 50 µL of InstaGene Matrix (BioRad Laboratories, Hertfordshire, UK) plus a small quantity (approximately 8 mg) of sterile silica powder to ease egg tissue and cell disruption. Then, the content of each tube was mashed with a sterile pestle and processed for DNA extraction following the manufacturer’s instructions. DNA extraction was also performed from liquid samples of the water or rinse water from treated eggs: 1.5 mL of CW, 1.5 mL of PAW and 1.5 mL of SHTW, were transferred in Eppendorf tubes and centrifuged at 13,000 rpm for 8 min. The supernatant of each sample was replaced by 25 μL of InstaGene Matrix and processed for DNA extraction following the manufacturer’s instructions. Finally, the supernatant of each vial (containing DNA from eggs or liquids) was transferred into another 1.5 mL tube and preserved at –20 °C until the molecular analyses. According to the DNA extraction, a DGGE analysis was performed to determine the presence of *Ca*. E. dacicola in the DGGE bacterial profiles before performing real-time PCR. Amplification of the V6-V8 region of the 16S rRNA gene was carried out with the universal primer pair 986F-GC and 1401R [43] in a 25-µL mixture containing 2 µL of template DNA, 1.5 mmol L^−1^ MgCl_2_, 200 mmol L^−1^ of each deoxynucleotide triphosphate (dNTP) (Promega Corporation), 10 pmol of each primer (TIB MolBiol), 1x green GoTaq^®^ flexi buffer (Promega), and 1 U of GoTaq^®^ polymerase (Promega). The reaction conditions were as follows: 94 °C for 4 min, followed by 35 cycles of denaturation at 95 °C for 45 s, annealing at 55 °C for 45 s, and extension at 72 °C for 45 s; and final extension at 72 °C for 7 min. Three independent PCR amplifications were performed for each sample, and the triplicate amplification products were pooled to minimize the effect of PCR biases. The amplification products were loaded onto a polyacrylamide gel (acrylamide/bis 37.5:1; Euroclone), with a linear denaturing gradient obtained with a 100% denaturing solution containing 40% formamide (Euroclone) and 7 M Urea (Euroclone). The gels were run for 17 hours in 1X TAE buffer at constant voltage (80 V) and temperature (60 °C) using the INGENY phorU-2 System (Ingeny International BV). Then, gels were stained with SYBR^®^GOLD (Molecular Probes) diluted 1:1,000 in 1X TAE, and the gel images were digitized using a Chemidoc XRS apparatus (Bio-Rad).

### DNA extraction from flies

*B. oleae* flies were killed by freezing at –20 °C for 15 min, washed with a 2% sodium hypochlorite solution and then rinsed twice in deionized sterile water in a laminar flow hood. Each adult’s head was dissected under a stereoscopic microscope with sterile tools, and the oesophageal bulb was extracted. DNA extraction of each bulb was carried out as described above for eggs. DNA extracted from the oesophageal bulbs of wild *B. oleae* flies was amplified as described above and used as a *Ca*. E. dacicola positive control in end-point PCR and as a marker in DGGE analysis, and it was used to construct the standard curve for the real-time PCR. DNA was also extracted from the oesophageal bulbs of *B. oleae* flies developed from eggs than had been externally treated with the SHTX mixture. Amplification followed by DGGE was performed as described above.

### Real-time PCR

Quantitative real-time PCR analysis was performed with primers EdF1 [23] and EdEnRev [44] was used to determine the relative abundance of *Ca*. E. dacicola varied across eggs surface treatments. Amplifications were carried out using a CFX96 Real-Time PCR Detection System (Bio-Rad Laboratories, Hertfordshire, UK) in a 20-μL mixture containing 2X SsoAdvanced Universal SYBR^®^ Green Supermix (Bio-Rad), 400 nmol/L of each primer and 2 μL of template DNA. The amplification conditions involved denaturation at 95 °C for 3 min, followed by 40 cycles of 95 °C for 15 s and 60 °C for 30 s. Fluorescence data were collected at the end of the hybridization step. Amplicon specificity was tested with a dissociation curve analysis by increasing the temperature by 0.5°C every 30 s from 65 to 95 °C. Negative controls and standard curves were run on each plate. The standard curve was prepared with a sample of DNA extracted from the bulb of a wild *B. oleae* female with *Ca*.E. dacicola and 5-fold serially diluted. The efficiency of the primer pair (E) was determined by calculating the slope of the log-scale standard curve and applying the following equation: E=10^(−1/slope)^ [45]. Each standard dilution and unknown sample was run in triplicate, and the threshold cycle (Ct) of these technical replicates were averaged for each individual sampled. The relative abundance of *Ca*. E. dacicola (R) was calculated according to Estes et al. [42]. The number of copies of *Ca*.E. dacicola 16S rRNA gene in egg samples treated with sodium hypochlorite (SHTXE) or propionic acid (PAE) or in water samples where eggs had been taken (CW, PAW, SHTXW) (E_sample_) was normalized relative to the number of copies of *Ca*.E. dacicola 16S rRNA gene found in egg samples washed with water (E_CE_) according to the formula: 
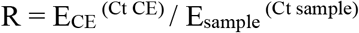

Four separate real-time PCR amplifications were performed using egg samples from four experimental replicates conducted over time, and the data from each treatment were averaged over the four replicates. Quantitative real-time PCR analysis was also performed with universal primers 338F-518R [46], as described above, to determine the relative abundance of bacteria on eggs surface and rinse water as well as.

### Sequence analysis

The middle portions of several DGGE bands were aseptically excised from the gel and directly sequenced by Macrogen Service (Macrogen LTD, The Netherlands). The sequence chromatograms were edited using Chromas Lite software (v.2.1.1; Technelysium Pty Ltd.; http://www.technelysium.com.au/chromas-lite.htm) to verify the absence of ambiguous peaks and to convert them to FASTA format, DECIPHER’s Find Chimeras web tool (http://decipher.cee.wisc.edu) was used to uncover chimeras in the 16S rRNA gene sequences. The sequences were analyzed via the web-based BLASTN tool (NCBI; http://www.ncbi.nlm.nih.gov/BLAST) of GenBank to identify bacterial species of highest similarity. The nucleotide sequences were deposited in the GenBank database under accession numbers MG800838 to MG800842.

### Scanning electron microscopy

Fifty eggs of each treatment were dehydrated in a series of graded ethanol from 50% to 99%, with 15 min at each grade. After dehydration, the eggs were allowed to dry under a hood at room conditions. On each aluminum stub, at least 5 eggs were mounted, taking care to arrange them horizontally to obtain a clear view of the area underlying the micropylar cup, which corresponds to the base of the egg anterior pole. Mounted eggs were gold-sputtered using a Balzers Union^®^ SCD 040 unit (Balzers, Vaduz, Liechtenstein). For the observations carried out at the Electronic Microscopy Labs at SIMAU, Polytechnic University of Marche, a FE-SEM Zeiss^®^ SUPRA 40 scanning electron microscope (Carl Zeiss NTS GmbH, Oberkochen, Germany) and a Philips^®^ XL 30 scanning electron microscope (Eindhoven, The Netherlands) were used. Additional investigations were conducted at the Department of Agricultural, Food and Agro-Environmental Sciences, University of Pisa, using a FEI Quanta 200 high-vacuum scanning electron microscope. The densities of the bacterial colonies present on the eggs from the three treatments were determined by counting the number of visible rods in a sample area enclosed by an electronic rectangular frame (approximately 800 *μ*m^2^) applied to the SEM screen where the base of the egg anterior pole was visible.

### Statistical analyses

Quantitative data from real-time PCR and data on the bacterial colonies on the egg surface (after square-root transformation to satisfy normality requirements) were analyzed through one-way analysis of variance (ANOVA) followed by Tukey’s honestly significant difference (HSD) test for means separation (P≤0.05) [47]. All of the analyses were performed using Statistica 6.0 (Statsoft, Italy).

## Results

### DGGE analysis

The first experiment was conducted to detect the presence of *Ca*. E. dacicola on the surface of *B. oleae* eggs. The PCR-DGGE profiles of egg samples washed with water (CE) showed more complex band patterns than did those obtained from egg samples treated with propionic acid (PAE) and the mixture hypochlorite + TritonX (SHTXE) or samples of water CW, PAW and SHTXW (Fig. 1). In each DGGE profile of eggs treated with water, a clear band was consistently present that showed the same migration behavior as the band formed by the sample of the oesophageal bulb of *B. oleae* used as marker of *Ca*. E. dacicola (M). This band was also present in the other DGGE profiles and showed a decreasing intensity from CE > PAE > SHTXE and rinse water samples.

**Figure 1.**
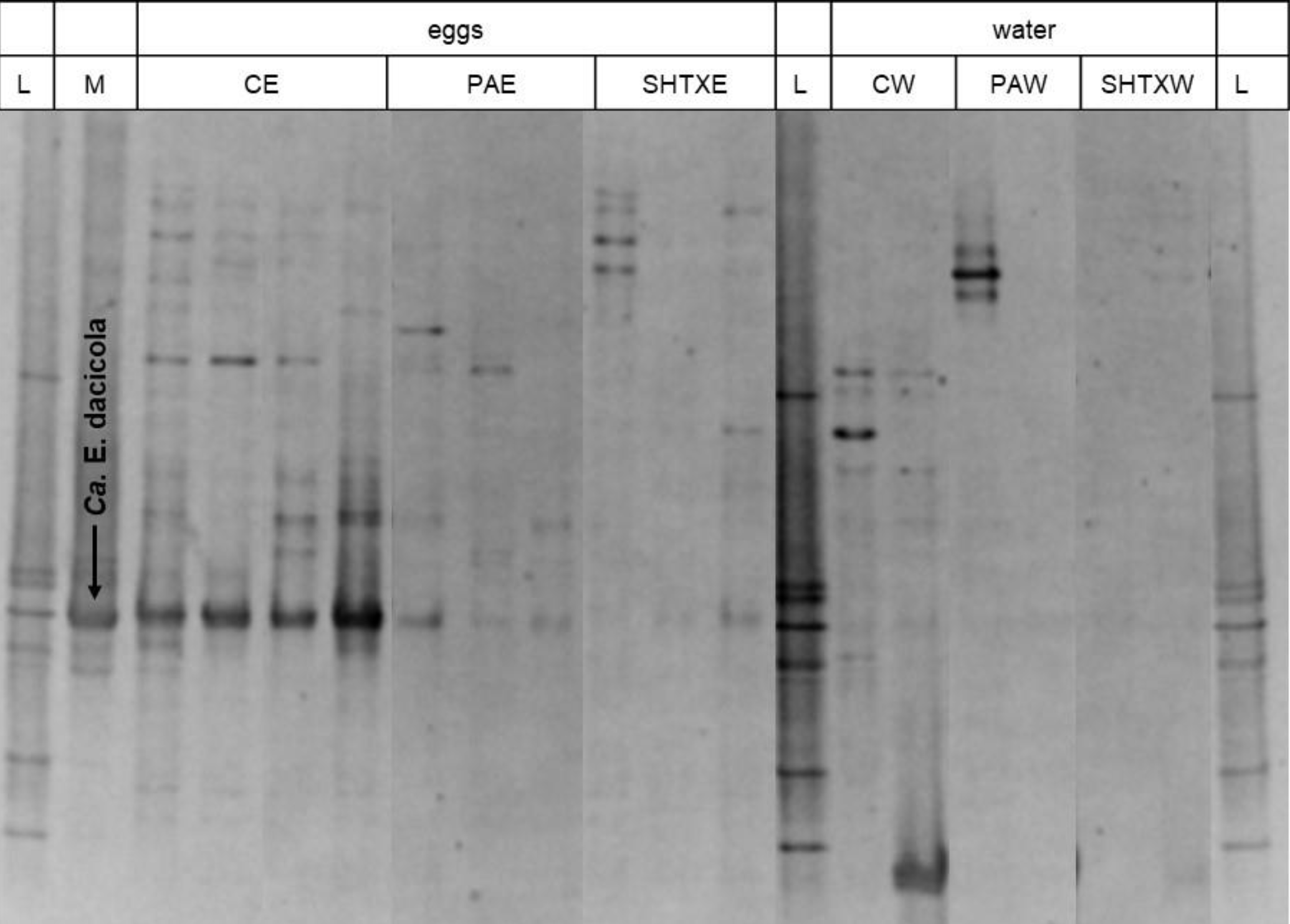
PCR-DGGE profiles of the 16S rRNA gene fragments obtained by amplification of DNA extracted from egg samples and rinse water. DGGE denaturing gradient 42–68%. Arrowed band indicates a DNA fragment obtained by amplification of DNA extracted from wild fly oesophageal bulbs and used as species marker of *Ca*. E. dacicola. L, ladder; M, 16S rRNA gene fragment obtained by amplification of DNA extracted from the oesophageal bulb of a wild fly and used as marker of *Ca*. Erwinia dacicola; CE, eggs washed with water (control eggs); PAE, eggs treated with 0.3% propionic acid; SHTXE, eggs treated with sodium hypochlorite + Triton X mixture; CW, water from control eggs; PAW, second rinse water after treatment with PA; SHTXW, second rinse water after treatment with SHTX.

### Relative abundance of *Ca*. E. dacicola in *B. oleae* eggs

The analysis of the presence of *Ca*. E. dacicola on *B. oleae* eggs laid by wild females and treated with disinfectants showed that the amount of the symbiont was decreased in the eggs of the various treatments relative to eggs of the control treatment (Fig. 2). Specifically, the quantity of the symbiont was reduced nearly by 2 times in eggs handled with the propionic acid solution (0.503±0.066 relative abundance of *Ca*. E. dacicola in PAE *vs Ca*. E. dacicola in CE), whereas in SHTXE, the bacterial load was decreased by approximately 5 times (0.211±0.125 relative abundance of *Ca*. E. dacicola in SHTXE *vs Ca*. E. dacicola in CE) relative to the quantity in the CE. One-way ANOVA revealed significant differences among the treatments (F_2,9_ = 95, *p*<0.001), and post hoc HSD tests revealed significant differences between the various treatments and the control treatment.

**Figure 2.**
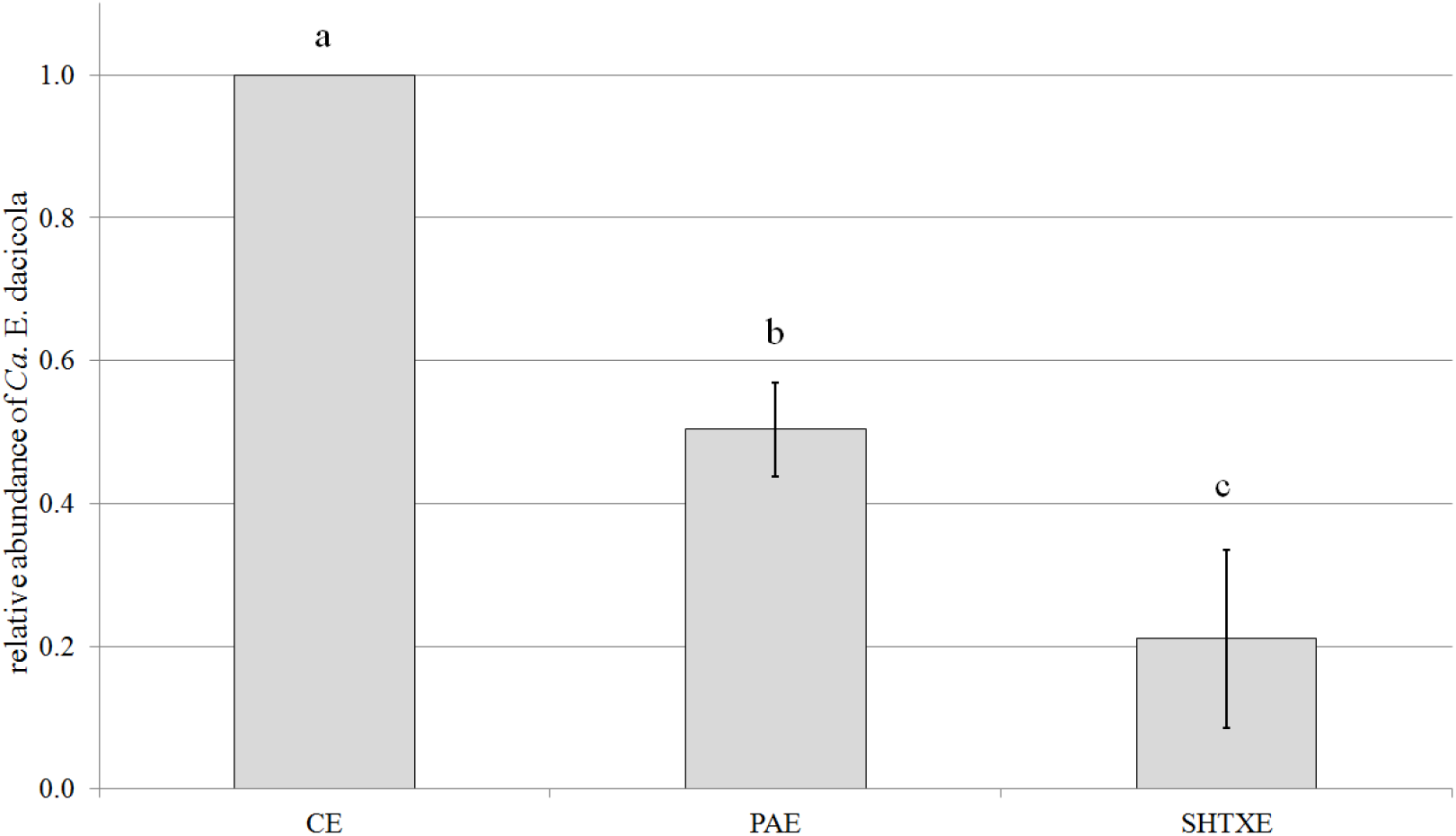
Relative abundance of *Ca*. E. dacicola (mean ± SD) in eggs washed with water (CE, control eggs) considered equal to 1 in comparison with eggs treated with 0.3% propionic acid solution (PAE), or with sodium hypochlorite + Triton X (SHTXE). One-way ANOVA followed by Tukey’s test at P≤0.05 (n=4) was performed; different letters above bars indicate significant differences between treatments.

Real-time PCR was performed on the rinse water of the three treatments to evaluate *Ca*. E. dacicola presence (Fig. 3). As expected, the relative abundance of the symbiont in the two rinse waters PAW and SHTXW was very low (0.00109±0.00017 and 0.0003±0.00021 relative abundance of *Ca*. E. dacicola in PAW and SHTXW, respectively, *vs Ca*. E. dacicola in CE). The water CW contained a greater quantity of *Ca*. E. dacicola (0.2349±0.31225 relative abundance of *Ca*. E. dacicola in CW *vs Ca*. E. dacicola in CE). Statistically significant differences were detected among treatments, with the bacterial content of the control rinse water comparable to the bacterial load on the eggs treated with both disinfectants (F_2,15_ = 59 M, *p*<0.001). However, considerable amounts of the *B. oleae* symbiont are lost even when eggs are washed with water; the load was assessed via real-time PCR analysis as representing approximately 20% of the original load.

**Figure 3.**
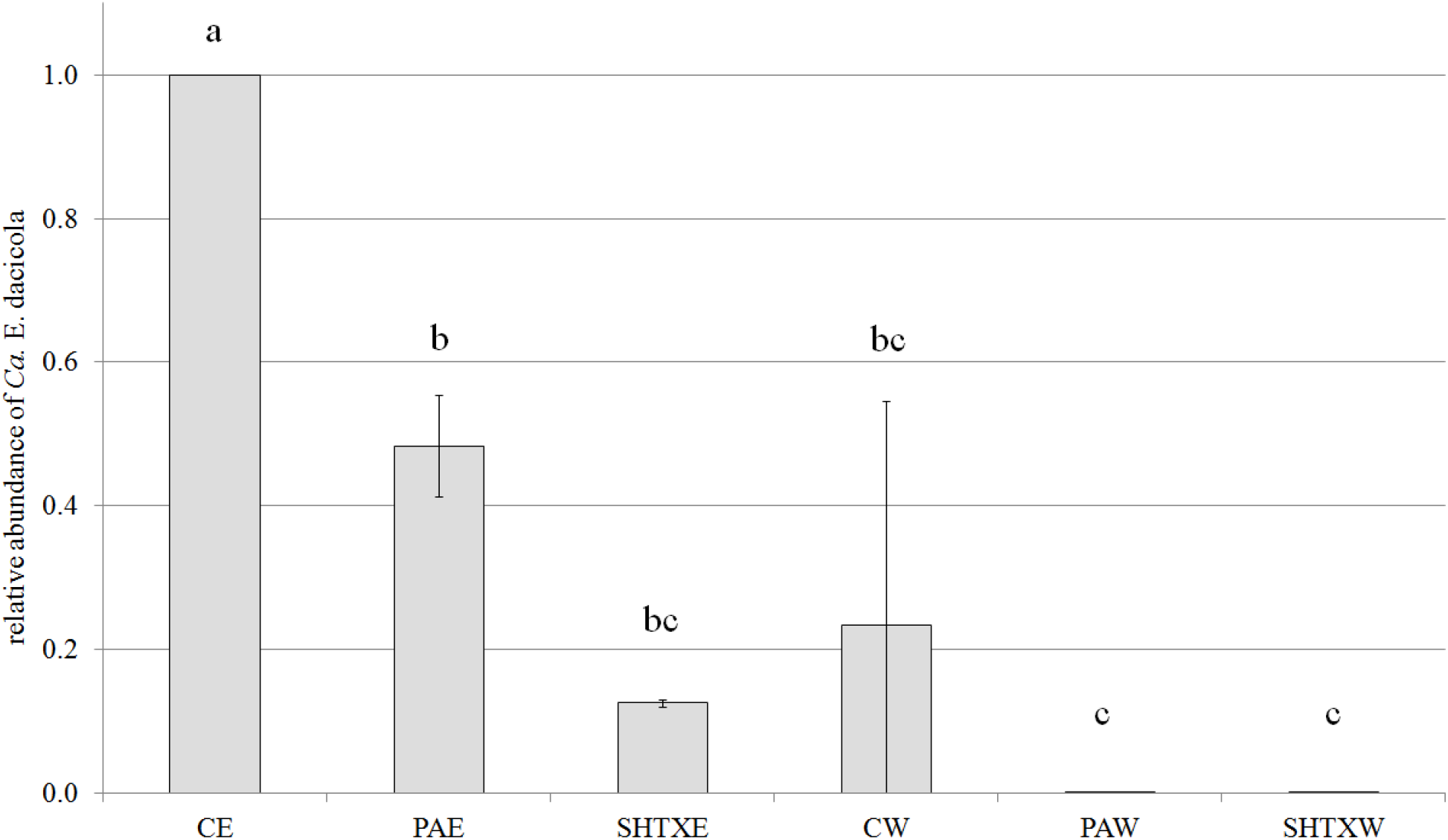
Relative abundance of *Ca*. E. dacicola (mean ± SD) in eggs washed with water (CE, control eggs) considered equal to 1 in comparison with eggs treated with 0.3% propionic acid solution (PAE), sodium hypochlorite + Triton X (SHTXE) and the respective rinse water CW, PAW, SHTXW. One-way ANOVA followed by Tukey’s test at P≤0.05 (n=3) was performed; different letters above bars indicate significant differences between treatments.

### Morphological observations

Eggs treated with the two disinfectants (PAE and SHTXE) or washed only with water (CE) were observed via SEM. The egg of *B. oleae* is elongated and slightly curved (whole egg not shown); it is characterized by a well-developed anterior pole with an overturned cup-like protrusion that is supported by a short peduncle, forming the micropylar apparatus (Fig. 4A and 4C). The protrusion margins display several knobs forming a festooned rim, which give the micropylar apparatus the overall appearance of a balloon tuft. The micropylar aperture is located in the center of the protrusion, and the peduncle shows several large openings connected with internal chambers (Fig. 4). Eggs washed with water showed many rod-shaped bacterial colonies scattered on the micropylar apparatus as well as on its base, around the openings of the internal cavities (Fig. 4B). In contrast, all the samples of eggs treated with SHTX or PA showed a total lack or negligible quantity of bacterial masses on the chorionic surface of the anterior pole (Fig. 4A, 4C, 4D). Counts of the number of bacterial colonies within an electronic frame confirmed that treatment with the disinfectants greatly affected the presence of bacteria (F_2,12_ = 23.57, *p*<0.001). PAE and SHTXE showed significant reductions of bacterial colonies relative to the colonies on CE (Fig. 5).

**Figure 4.**
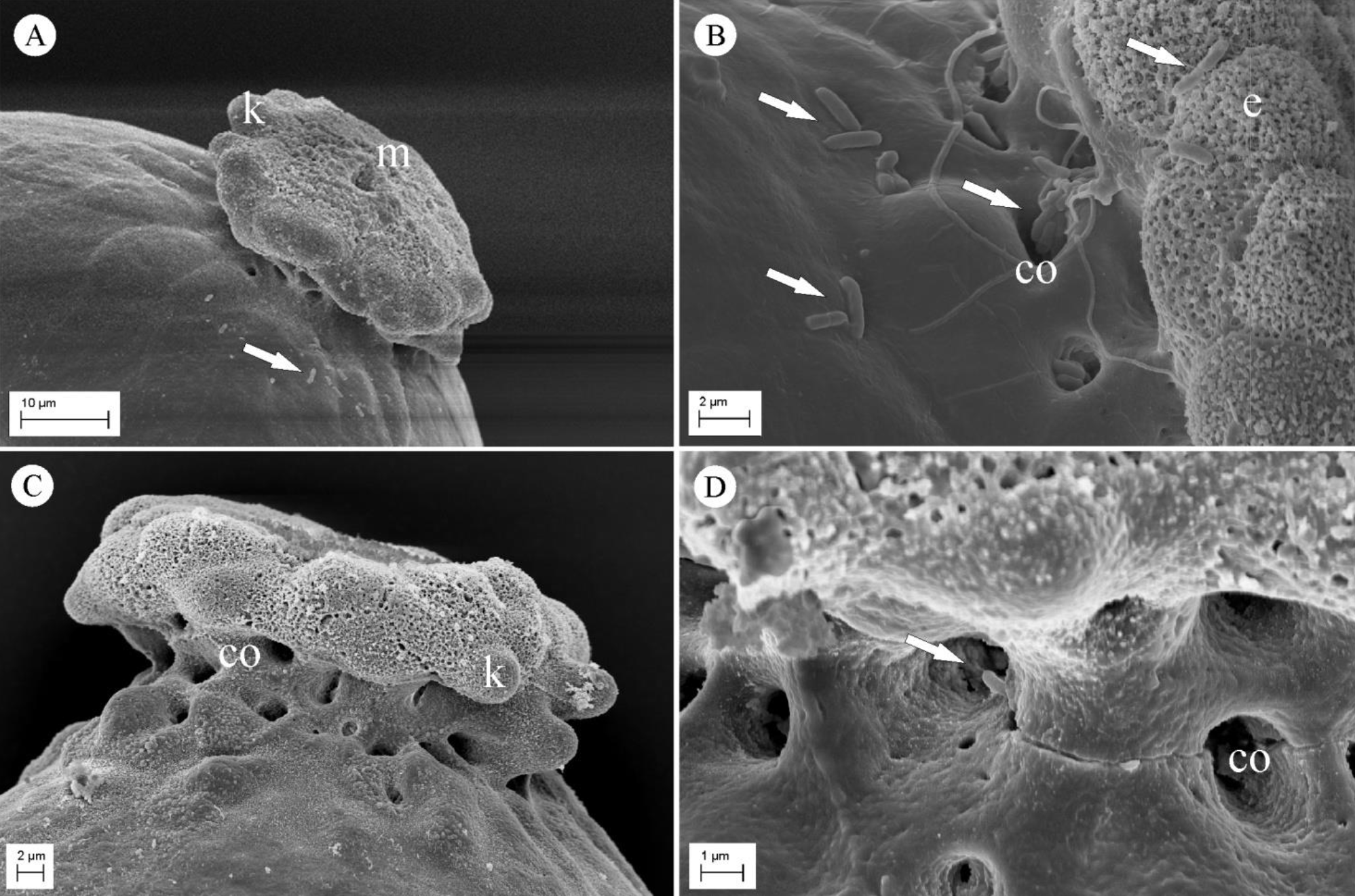
Scanning electron micrographs of the anterior pole of *B. oleae* eggs. (A) Anterior pole of an egg treated with 0.3% propionic acid showing the reduction in the number of bacterial cells on the egg surface. (B) Magnification of an egg washed with water (control) showing the bacterial cells scattered on the micropylar apparatus and around the openings of the internal cavities. (C) Anterior pole of an egg treated with sodium hypochlorite + Triton X mixture (SHTX) showing the absence of bacteria on the egg surface. (D) Magnification of the base of the micropylar apparatus of an egg treated with sodium hypochlorite + Triton X mixture (SHTX) displaying a single bacterial cell (arrow) in an internal cavity opening. Arrows indicate rod-shaped bacteria; (co) cavity opening; (e) exochorionic layer with characteristic sponge-like feature; (k) knobs on protrusion margins; (m) micropylar opening.

**Figure 5.**
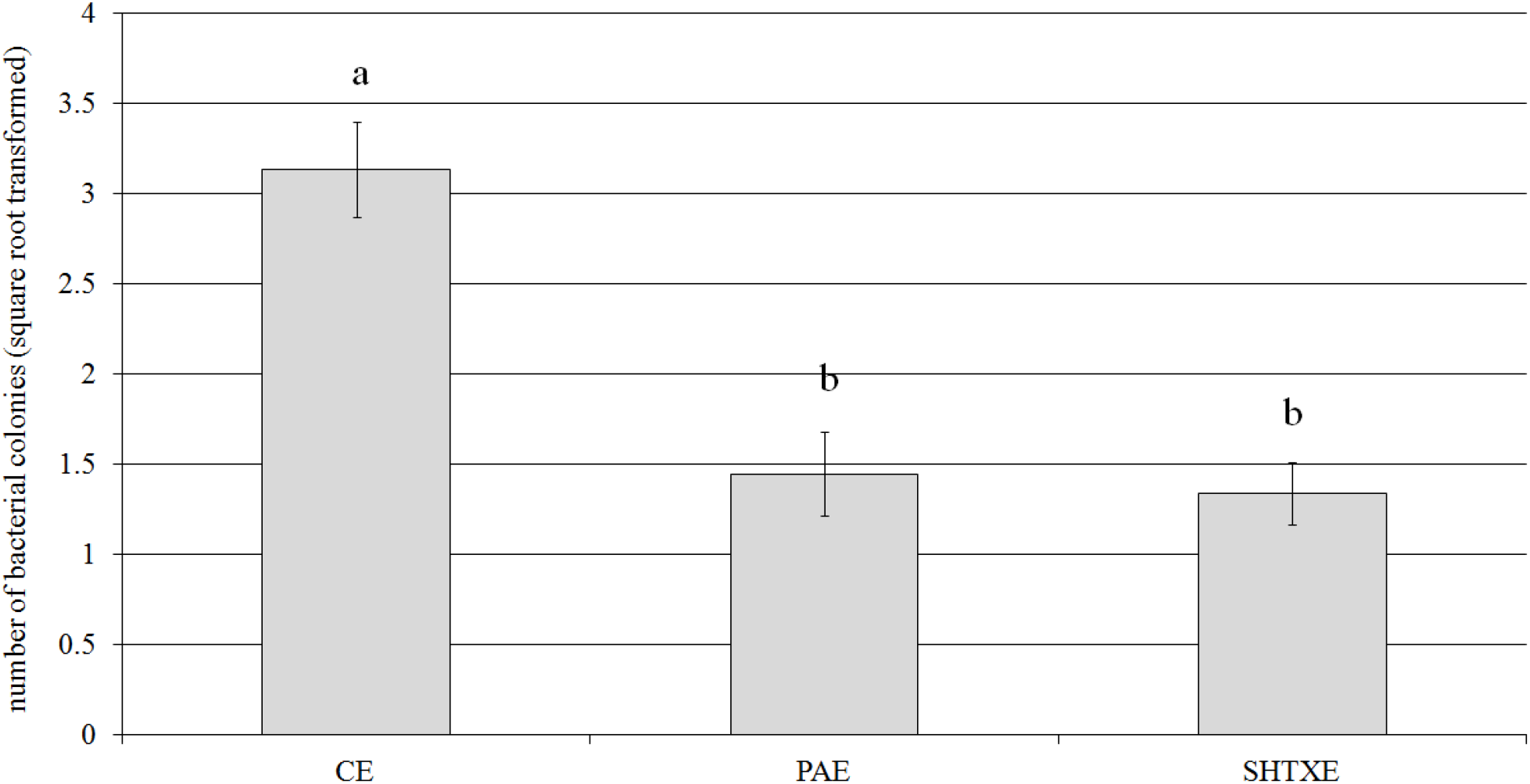
Number of bacteria (mean ± SD) counted within an electronic frame in the area close to the cup-like protrusion of *B. oleae* eggs washed with water (CE) or after treatment with 0.3% propionic acid solution (PAE) or sodium hypochlorite + Triton X mixture (SHTXE). One-way ANOVA followed by Tukey’s test at P≤0.05 (n=5) was performed; different letters above bars indicate significant differences between treatments.

**Figure 6.**
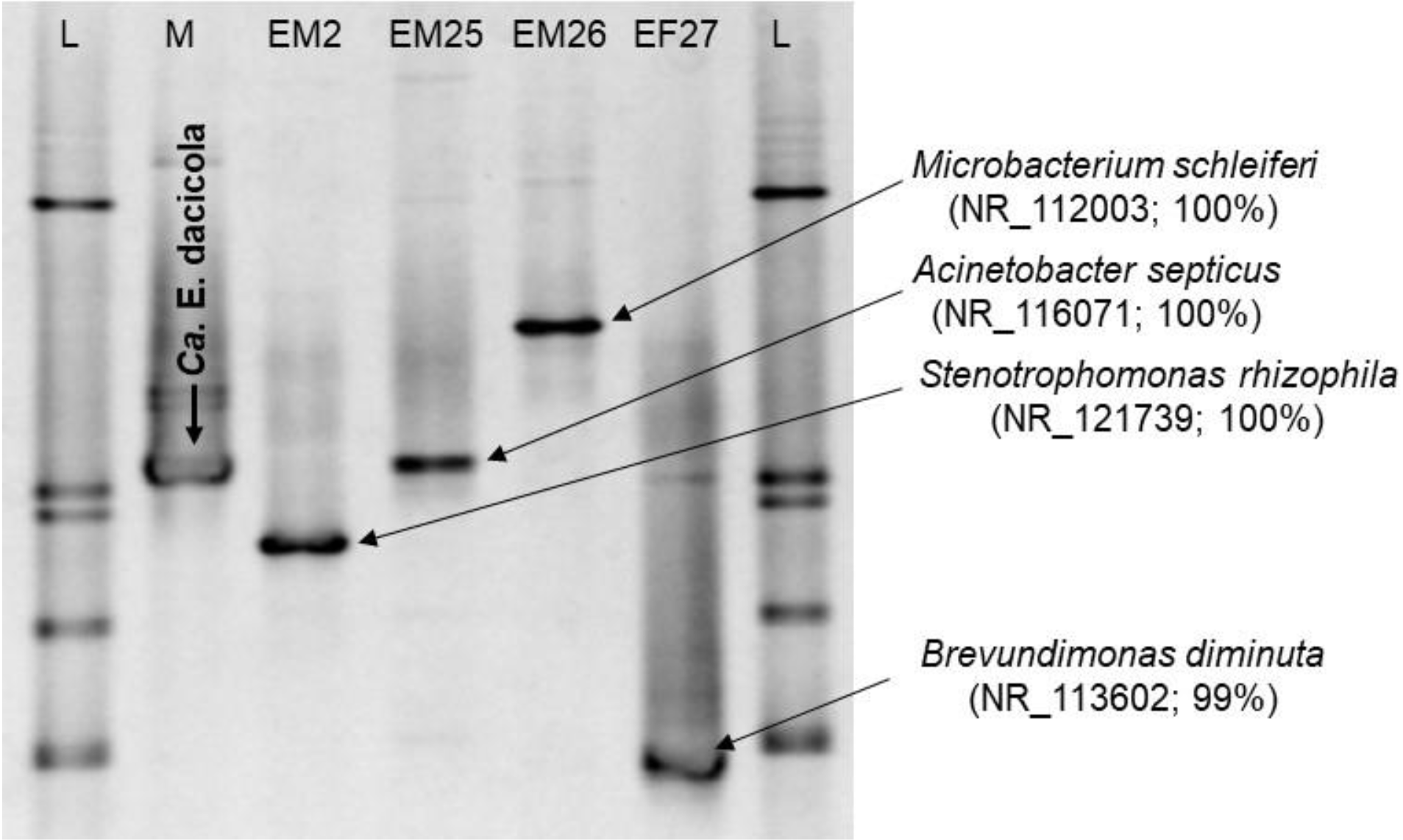
PCR-DGGE profiles of the 16S rRNA gene fragments obtained by amplification of DNA extracted from the oesophageal bulb of wild *B. oleae* flies and *B. oleae* flies developed from eggs externally treated with SHTX (1% sodium hypochlorite + 0.1% Triton X mixture). DGGE denaturing gradient 48–65%. Arrowed bands indicate band excised; GenBank accession number and % sequence similarity of the nearest BLAST match are also reported. L, ladder; M, 16S rRNA gene fragment obtained by amplification of DNA extracted from the oesophageal bulb of a wild fly and used as marker of *Ca*. Erwinia dacicola; EM2, EM25–27, sample codes.

### Progeny development

Egg hatchability was low and did not differ among the treatments: on average, it was 35.99±8.01% for CE, 34.29±7.13% for PAE and 36.64±21.11% for SHTXE (4 replications; the number of eggs per treatment varied from approximately 30 to 100). Moreover, the pupal recovery was very low and variable among treatments: 6.43% (from 184 eggs) for CE, 3.42% (from 147 eggs) for PAE and 13.56% (from 189 eggs) for SHTXE (percentages from the pooled data of 3 replications). Ultimately, only a few adults per treatment emerged from pupae reared on artificial diet: 11 from CE, 5 from PAE and 11 from SHTXE. A positive amplification product was obtained only from four oesophageal bulbs of flies that developed from SHTXE and their PCR-DGGE profiles are reported in Fig. 6. Each amplicon showed a characteristic migration pattern that differed from that produced by the *Ca*. E. dacicola marker. Bands were removed from the DGGE gels and sequenced, revealing their similarities to *Stenotrophomonas rhizophila* (100% similarity to GenBank accession number NR_121739), *Microbacterium schleiferi* (100% similarity to GenBank accession number NR_112003), *Brevundimonas diminuta* (99% similarity to GenBank accession number NR_113602) and *Acinetobacter septicus* (100% similarity to GenBank accession number NR_116071).

## Discussion

The main objective of this research was to evaluate the impact of disinfectants on the presence of *Ca*. E. dacicola on *B. oleae* eggs that had been laid by wild females. The goal of our research program is to establish a symbiotic olive fruit fly strain comprised of vigorous adults and with stable, high performance to produce males that can compete with wild specimens in sterile insect technique applications. Our findings show that only those eggs washed with water (CE) maintained most of the bacterial load delivered by the mother to the egg surface during oviposition. The bacterial symbiont on the collected eggs was *Ca*. E. dacicola, as evidenced by PCR-DGGE analysis, confirming previous studies [41].

According to our real-time PCR and SEM observations, eggs treated with PA, the antifungal agent recommended as part of standard olive fruit fly rearing procedures [32, 48], can lose up to half of the content of the symbiont transferred by the mother. Propionic acid was first evaluated and selected from among several disinfectants for its non-negative effects on egg hatching in the 1970s, when rearing procedures of the olive fruit fly were first established [49]. Propionic acid and propionates are considered as “Generally Recognized As Safe” (GRAS) food preservatives for humans. They are used as mold inhibitors and disrupt proton exchange across membranes, thereby negatively affecting amino acid transport [50]. In insect rearing protocols, propionic acid solutions are commonly recommended and used as antifungal agents, but they are considered ineffective against bacteria [51, 52]. It is likely that in our experiments, PA treatment significantly reduced the symbiont presence by facilitating the mechanical removal of bacteria from the egg surface during egg washing. Regardless of the mechanism, we can conclude that its usage eliminates most of the *Ca*. E. dacicola cells transferred from the mothers to their eggs.

The second washing treatment used in our experiment was a mixture containing sodium hypochlorite and Triton X (SHTX). This mixture was used to obtain results that can be compared to those obtained by Estes et al. [42]. Sodium hypochlorite is widely used at mild concentrations to surface-sterilize insect adults before dissection, but it is also recommended for the surface sterilization of eggs for insect rearing [53]. Since bleach is a very effective bactericide, we expected a severe reduction of *Ca*. E. dacicola following the treatment of *B. oleae* eggs with the treatment mixture. Moreover, some of the bacteria present on the egg surfaces were likely removed by the combined surfactant action of Triton X. A detectable quantity of other bacteria, as evidenced by amplification with universal primers, was observed only for the control water (CW) (data not shown). Exposure of DNA to sodium hypochlorite causes cleavages in DNA strands, breaking the DNA into small fragment or individual bases that precluded its amplification [54]. Therefore, we hypothesize that both PA and SHTX destroyed bacterial DNA, precluding the 16S rRNA gene amplification in rinse water.

Considering our findings along with those of Estes et al. [42], we can better understand the importance of avoiding the loss of the symbiont from eggs. The relative abundance of *Ca*. E. dacicola in eggs laid by wild females had been estimated as being approximately 5,000 times lower than that in the larval stage [42]. Furthermore, the symbiont can grow and colonize the gastric caeca in the larval midgut. Thus, we speculate that common lab rearing procedures may reduce or remove the bacterial load under a minimum threshold symbiont egg load necessary to maintain the symbiotic relationship. We believe that to prevent reductions in bacterial transmission, efforts should be made to avoid the usage of disinfectants in egg collection and/or to establish an oviposition substrate like olives where females can directly oviposit, as has been attempted with various fruits [55, 56].

It is generally known that common procedures used in lab rearing can affect the presence of microorganisms that are associated with insects in complex symbioses. The importance of the gut microbiota in the mass rearing of the olive fruit fly has been recently noted, and new rearing methods and diets have been recommended [31, 57].

When insects are reared in a laboratory, small-scale insectary or large-scale facility, they are exposed to several sources of contamination, which are enhanced by diverse factors such as the artificial, constrained environment; the non-natural diet; and the high population density [53, 58]. For this reason, various antimicrobials are used to prevent the growth of potentially harmful microorganisms (pathogenic or non-pathogenic contaminants) in different phases of the rearing process [52, 58]. The current procedure used to rear the olive fruit fly [48] was established after numerous experimental tests to evaluate several technical conditions as well as all diet ingredients; however, the maintenance of the bacterial symbiont in the insect colony was not considered. Moreover, lab populations of the olive fruit fly, reared for successive generations under artificial conditions, have shown deleterious biological, genetic and behavioral changes [59, 60, 61]. Such alterations might be due to different causes, and antimicrobials and antibiotics are likely to be important modifying agents. Streptomycin has been shown to negatively affect *B. oleae* larval growth [62], and nipagin has been shown to change the fly’s microflora composition, causing variations in Adh allele frequencies [63]. Fitness reductions caused by antimicrobial agents have been documented in other insects, such as members of Hemiptera [64] and Lepidoptera [65]. Taking into consideration recent findings on the olive fruit fly endosymbiont, *Ca*. E. dacicola, the indirect effects of piperacillin on adult fitness in *B. oleae* have been evaluated [28]. In addition, the toxicity of the different disinfectants used in artificial larval diets should be tested for potential destructive effects on the symbiont.

It is believed that bacterial symbionts are transmitted from olive fruit fly females to the progeny via eggs: this process was hypothesized by Petri [20, 21] and well documented by Mazzini and Vita [30]. Through SEM and TEM observations, these latter authors described the ovarian eggs and female reproductive organs as being devoid of bacteria, whereas the rectal, finger-like diverticula that converge into the ovipositor base harbor many bacterial masses. However, bacterial colonies have since been found close to the anogenital opening of the olive fruit fly female [24]. The absence of bacteria in ovarian eggs was also confirmed [66] in a study of the structure and morphogenesis of the *B. oleae* egg shell and micropylar apparatus. Moreover, submicroscopic observations have confirmed the absence of bacteria inside the vitelline membrane and the occasional occurrences of bacteria in the micropylar canal [30]. Based on these previous investigations, we can state that newly hatched larvae acquire bacterial symbionts from the cavities that underlie the micropylar apparatus, where bacteria likely grow during olive fruit fly embryogenesis and where the larva mouthparts burst at egg eclosion [67]. Our observations revealed the presence of bacterial cells over and around the micropylar apparatus, with some cells occurring inside the cavity opening.

Further insight into the symbiont’s transfer can be drawn from the egg morphology of *B. oleae*. Based on previous studies [30, 66] and our SEM observations, we hypothesize that the peculiar morphology of the micropylar apparatus might be related to the transmission of the symbiont. The balloon tuft-like protrusion of the anterior pole appears to be a potentially advantageous structure for scraping bacteria from the lumen of the rectal tract, where the diverticula release their bacterial content. According to earlier studies [68] and our investigations, *B. oleae* eggs exit from ovaries with the posterior pole directed toward the ovipositor. In this way, eggs entering the ovipositor cross throughout the poky passage and are covered with bacteria that occur mainly around and below the protrusion of the micropylar apparatus. Eggs are then laid inside the olive, oblique to the surface and with the anterior pole close to the pierced fruit skin [69, unpublished observations of the authors). The egg morphology of different species belonging to or closely related to the *Bactrocera* genus has not received much attention: apart from some notes on *Zeugodacus cucurbitae* (Coquillet) and *B. dorsalis* (Hendel) [70], only one research, carried out using SEM, investigated the eggs of *B. carambolae* Drew and Hancock and *B. papayae* Drew and Hancock [71], the latter, recently synonymized to *B. dorsalis* [72]. None of these species display the characteristic shape of the anterior pole of *B. oleae* egg. Furthermore, eggs of *Anastrepha* species, which have been thoroughly studied, have a different micropylar shape [73]. Thus, it would be interesting to analyze and compare the micropylar structures of different species with reference to symbiont transmission. Our initial findings on the development of eggs treated with antimicrobials appear to suggest that different bacteria may settle in the oesophageal bulb after the removal of most of the bacterial load from the eggs, including the symbiont load, as occurred after washing the eggs with SHTX. The four bacterial species recovered from flies are very different: *Stenotrophomonas, Brevundimonas* and *Acinetobacter* are genera of gammaproteobacteria belonging to the Pseudomonadales order, whereas *Microbacterium* is a genus of Actinobacteria. These species may be considered ubiquitous.

*M. schleiferi* and *S. rhizophila* have been isolated from air, soil, water, and plants as well as from larval and insect guts [74]. *B. diminuta* is considered a major actor in the process of tissue decomposition as one of the most common organisms in the soil and other moist environments [75]. Isolates of *Brevundimonas vesicularis* were retrieved from the oesophageal bulb of wild olive flies using culture-dependent techniques in a survey aimed at studying the microbial ecology of *B. oleae* in Tuscany [33]. Although ubiquitous, *A. septicus* has mainly been isolated from animal and insect specimens (for example, *Anopheles gambiae*) and nosocomial infections [76].

Finally, considering that 1) we demonstrated a negative effect of disinfectants on the olive fruit fly symbiont, 2) olive flies can be reared on artificial diet without antibiotics for eight generations [41], 3) genetic changes can be avoided by refreshing lab colonies every five to eight generations with wild flies [36], and 4) *Ca*. E. dacicola can be transferred horizontally among adults through cohabitation, as recently shown [26], we assert that a stable symbiotic strain of the olive fruit fly can be established and maintained under lab conditions.

## Conclusions

Wild populations of the olive fruit fly benefit from the symbiont *Ca*. E. dacicola in the larval and adult stages, while lab colonies, which lack the symbiont, display reduced fitness. However, SIT applications rely on the availability of high-quality, mass-reared insects. To establish a symbiotic laboratory strain of the olive fruit fly, *Ca*. E. dacicola must be maintained in all of the fly’s developmental stages to produce high performing males and females. This research demonstrated that common disinfectants and antimicrobials used in egg collection strongly affect symbiont transmission from mother to progeny, with severe consequences, especially considering the bacterial “bottleneck” that naturally occurs in the transfer from female to larvae via the eggs. Negative effects of antibiotics on both wild and lab populations have been documented. This study demonstrated a direct detrimental effect of disinfectants commonly used in olive fruit fly rearing on *Ca*. E. dacicola. To maintain the bacterial-insect symbiotic relationship in lab strains, “it is crucial to provide rearing conditions that allow the normal maintenance of the interaction”, as Cohen stated [52]. Future research is needed to test different compounds and conditions for compatibility with symbiont presence in olive fruit fly lab colonies, especially during larval rearing using artificial diets, in which molds must be prevented. The findings of this research can be considered as a starting point for a general review of the entire rearing process for *B. oleae*.

## Author contributions

All of the authors conceived of and designed the experiments. PS, GB and RG reared the insects and performed the experiments. PS, GB and RG carried out insect dissection and DNA extraction. RP, GB and CV designed and performed the molecular biology techniques. SR and AB conducted the SEM observations. PS and RP analyzed the data. PS, RP, GB and AB wrote the manuscript. All of the authors participated in the revision of the initial draft and approved the final manuscript.

## Acknowledgements

The authors are grateful to the farm “Tenute Librandi Pasquale”, Vaccarizzo Albanese (Italy) and to Dr. Michele Librandi for providing olive fruit fly pupae used in the research. Thanks are also to Gabriele Ferrini for his valuable help in laboratory analyses. Antonio Belcari and Patrizia Sacchetti acknowledge the International Atomic Energy Agency of Vienna, Austria, for supporting their participation in the CRP “Use of Symbiotic Bacteria to Reduce Mass-Rearing Costs and Increase Mating Success in Selected Fruit Pests in Support of SIT Application” (D41024).

## Funding

This research was partially funded by the University of Florence (Fondi d’Ateneo) and by the “Fondazione Cassa di Risparmio di Firenze”.

## Availability of data and materials

The datasets used and/or analyzed during the current study are available from the corresponding author on reasonable request.

## Competing interests

The authors declare that they have no competing interests.

